# Boost in Test-Retest Reliability in Resting State fMRI with Predictive Modeling

**DOI:** 10.1101/796714

**Authors:** Aman Taxali, Mike Angstadt, Saige Rutherford, Chandra Sripada

## Abstract

Recent studies found low test-retest reliability in fMRI, raising serious concerns among researchers, but these studies mostly focused on reliability of individual fMRI features (e.g., individual connections in resting state connectivity maps). Meanwhile, neuroimaging researchers increasingly employ multivariate predictive models that aggregate information across a large number of features to predict outcomes of interest, but the test-retest reliability of predicted outcomes of these models has not previously been systematically studied. Here we apply ten predictive modeling methods to resting state connectivity maps from the Human Connectome Project dataset to predict 61 outcome variables. Compared to mean reliability of individual resting state connections, we find mean reliability of the predicted outcomes of predictive models is substantially higher for all ten modeling methods assessed. Moreover, improvement was consistently observed across all scanning and processing choices (i.e., scan lengths, censoring thresholds, volume-versus surface-based processing). For the most reliable methods, reliability of predicted outcomes was mostly, though not exclusively, in the “good” range (above 0.60).

Finally, we identified three mechanisms that help to explain why predicted outcomes of predictive models have higher reliability than individual imaging features. We conclude that researchers can potentially achieve higher test-retest reliability by making greater use of predictive models.

## 1 Introduction

Recent studies report troublingly low test-retest reliability of functional magnetic resonance imaging (fMRI) metrics including resting state functional connectivity (Noble et al. 2019) and task activation (Elliott et al. 2020). These studies have attracted substantial attention from clinical and translational neuroscientists because adequate test-retest reliability of fMRI is critical for its use in individual-differences research. If differences in functional imaging features (e.g., connectivity, activation, or other features) across individuals are not stable across scanning sessions, that is, if values of these imaging features lack consistency and/or agreement across sessions, then these features cannot serve as a basis for constructing predictively useful objective markers (i.e., “biomarkers”) of traits of interest (Nunnally 1970).

The current literature examining reliability in fMRI has been mostly focused on the reliability of *single* imaging features: individual connections in resting state connectivity maps and individual voxels or regions of interest in task activation maps. Neuroimaging researchers studying individual-differences are, however, increasingly moving away from univariate tests performed separately on each imaging feature and are instead utilizing multivariate predictive models (Klöppel et al. 2012; Orru et al. 2012; Woo, Chang, et al. 2017; Scheinost et al. 2019) (hereafter “predictive models”). These methods aggregate information across thousands of distributed brain features, yielding a single overall “best guess” about the outcome of interest. Predictive models are now widespread in the field and have been used to predict a range of psychologically- and clinically-relevant outcomes including cognitive skills (Sripada, Rutherford, et al. 2019; Sripada et al. 2018 Jan 1; Dubois et al. 2018), pain ratings (Wager et al. 2013; Chang et al. 2015; Woo, Schmidt, et al. 2017), sustained attention (Kessler et al. 2016; Rosenberg et al. 2016), schizophrenia status (Watanabe et al. 2014), and depression subtype/treatment response (Drysdale et al. 2017), among many others (Sui et al. 2020). To date, however, the test-retest reliability of predicted outcomes derived from predictive models has not been evaluated.

This question is particularly interesting in light of well-known results from psychometrics that establish that aggregation of features, for example by taking sum scores or applying a weighting function, can yield a composite variable that is much more reliable than the individual items that make up the composite (Nunnally 1970; He 2009). Standard predictive models widely used in the imaging field work in just this way: they aggregate features through application of a weighting function (Woo, Chang, et al. 2017). Thus, it is possible that like composite scores in psychology, predicted outcomes from such models will exhibit meaningfully higher reliability than individual features.

To test this hypothesis, we turned to the Human Connectome Project (HCP) dataset (Van Essen et al. 2013), which has two sessions of resting state fMRI data acquired on consecutive days for a large sample of subjects. This dataset also has a large number of phenotypic outcome variables, allowing us to train predictive models across a number of psychological domains, including cognition, emotion, personality, and psychopathology symptom scores. We examined ten predictive modeling methods widely used in the neuroimaging field: Ridge Regression, Lasso, Elastic Net, Connectome Predictive Modeling (CPM) (Shen et al. 2017) (two versions), Brain Basis Set (BBS) (Sripada, Angstadt, et al. 2019) (two versions), Support Vector Regression (two versions), and Random Forest. Results from our systematic comparison showed that all predictive modeling methods yielded substantially higher test-retest reliability of predicted outcomes compared to individual connectivity features.

## 2 Methods

### 2.1 Subjects and Data Acquisition

All subjects and data were from the HCP-1200 release (Van Essen et al. 2013; WU-Minn HCP 2017). Study procedures were approved by the Washington University institutional review board, and all subjects provided informed consent. Four resting state runs were performed (14.5 minutes each run) across two days, with two runs the first day and two runs the second day. Data was acquired on a modified Siemens Skyra 3T scanner using multiband gradient-echo EPI (TR=720ms, TE=33ms, flip angle = 52°, multiband acceleration factor = 8, 2mm isotropic voxels, FOV = 208×180mm, 72 slices, alternating RL/LR phase encode direction). T1 weighted scans were acquired with 3D MPRAGE sequence (TR=2400ms, TE=2.14ms, TI=1000ms, flip angle = 8, 0.7mm isotropic voxels, FOV=224mm, 256 sagittal slices). T2 weighted scans were acquired with a Siemens SPACE sequence (TR=3200ms, TE=565ms, 0.7mm isotropic voxels, FOV=224mm, 256 sagittal slices).

### 2.2 Data Preprocessing

Processed volumetric and grayordinate data from the HCP minimal preprocessing pipeline that included ICA-FIX denoising were used. Full details of these steps can be found in Glasser (2013) and Salimi-Khorshidi (2014). Briefly, T1w and T2w data were corrected for gradient-nonlinearity and readout distortions, inhomogeneity corrected, and registered linearly and non-linearly to MNI space using FSL’s FLIRT and FNIRT. BOLD fMRI data were also gradient-nonlinearity distortion corrected, rigidly realigned to adjust for motion, fieldmap corrected, aligned to the structural images, and then registered to MNI space with the nonlinear warping calculated from the structural images. Then FIX was applied on the data to identify and remove motion and other artifacts in the timeseries. Volumetric images were smoothed with a 6mm FWHM Gaussian kernel.

The volumetric and grayordinate images then went through a number of resting state processing steps, including a motion artifact removal steps comparable to the type B (i.e., recommended) stream of Siegel et al. (2017). These steps include linear detrending, CompCor (Behzadi et al. 2007) to extract and regress out the top 5 principal components of white matter and CSF, high pass filtering at 0.008Hz. We evaluated the sensitivity of test-retest reliability to motion artifacts by employing two thresholds of excess framewise displacement for scrubbing frames: 0.5mm and 0.2mm.

### 2.3 Connectome Generation

We consider two standard atlases: i) for volumetric data, the parcellation of Power et al. with 264 ROIs of 4.24mm radius each (Power et al. 2011), ii) for grayordinate, the Gordon 333 parcel atlas (Gordon et al. 2016), augmented with parcels from high-resolution subcortical (Tian et al. 2020) and cerebellar (Diedrichsen et al. 2011) atlases. For each run, we calculated the Pearson’s correlation coefficients between the spatially averaged timeseries per ROI. These ROI-correlation matrices were then transformed using Fisher’s r to z-transformation and averaged across runs per day, yielding two connectivity matrices per subject, one for each day. We evaluated the effects of scan-length by utilizing ROI timeseries’ of different lengths for connectome generation. Keeping all 1200 points provided 14.5 minutes of data from each run (hereafter “29-minutes”); keeping the first 625 points provided 7.5 minutes of data from each run (hereafter “15-minutes”); and keeping the first 312 points provided 3.75 minutes of data from each run (hereafter “7.5-minutes”).

### 2.4 Inclusion/Exclusion Criteria

Subjects were eligible to be included if they had: 1) structural T1 data and had 4 complete resting state fMRI runs (14m 24s each); 2) full necessary behavioral data. To avoid confounding due to intra-familial similarity, we additionally randomly selected one individual from each sibship. This left 384 unrelated individuals to enter our main analysis.

### 2.5. ICC Metric for Test-Retest Reliability

Test-retest reliability assesses the consistency of scores measured across repeated sessions. Pearson product-moment correlations (*r*) are sometimes used as a reliability metric. A limitation of Pearson’s *r*, however, is that it only captures the similarity in the orderings between test-retest measurements (Cicchetti, 1994), so that if two sets of measurements are very far apart yet covary in the same order, they would be attributed high reliability. The intraclass correlation (ICC) statistic addresses this issue by centering and scaling the data using a pooled mean and standard deviation, similar to ANOVA (Cicchetti, 1994, Cicchetti, Sparrow 1981). There are several forms of ICC (Shrout and Fleiss). In the current study, imaging features are measured by the same “rater” (fMRI scanner) across two sessions, and we are interested in generalizing our results to repeated retest sessions. This situation corresponds to the ICC(2,1) statistic in the scheme of Shrout and Fleiss (1979), which we select as our ICC metric to assess the test-retest reliability of predicted outcomes of predictive models.

### 2.6. Types of Predictive Models

#### Lasso

Lasso is a form of regularized linear regression using an L_1_-norm penalty that tends to shrink coefficients to zero. In this way it can act as both a regularization and feature selection method. The amount of regularization is controlled via the lambda term in the objective function:

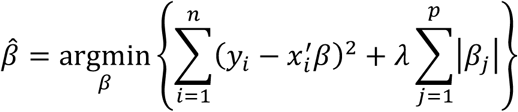

Lasso was performed using the glmnet package in Python (https://web.stanford.edu/∼hastie/glmnet_python/), and 100 automatically generated values of lambda were tested when fitting each model. On each fold of our 10-fold cross validation (see below), we used 5-fold cross validation within the training data to select the best lambda value.

#### Ridge Regression

Ridge regression (Hoerl and Kennard 1970) is another form of regularized linear regression is conceptually similar to Lasso. The main difference is that Ridge uses the L_2_-norm penalty, which constrains the sum of the squared coefficients. This results in shrinkage that is proportional to each coefficient’s magnitude, thereby eliminating the possibility that some coefficients with be regularized to zero. The amount of regularization is controlled by lambda in the objective function:

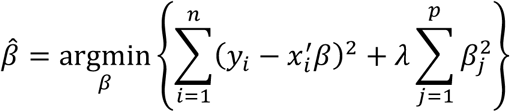

The Ridge Regression models were fit using the glmnet package in Python and 100 automatically generated values of lambda were tested when fitting each model. Lambda was chosen on each fold of our 10-fold cross validation (see below), using 5-fold cross validation within the training data.

#### Elastic Net

Elastic net (Zou and Hastie 2005) is a mix between an L_1_ penalized regression (i.e. lasso) and an L_2_ penalized regression (i.e. ridge regression). It attempts to balance the sometimes overly aggressive feature selection of lasso by mixing it with ridge regression. There is an additional hyperparameter, alpha, that controls the balance of the lasso and ridge penalties. The objective function for elastic net is:

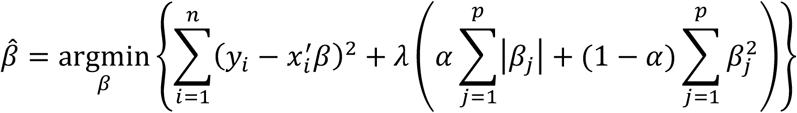

Elastic Net was performed using the glmnet package in Python; following Jollans and colleagues^32^, the alpha values were selected from a linear scale (0, 0.1, 0.325, 0.55, 0.775, 1) and the lambda values were selected from 100 automatically generated values. As before, within our overall 10-fold cross-validation (see below), we performed 5-fold cross-validation in the training data to select the best values for alpha and lambda.

#### Connectome Predictive Modeling (CPM)

Connectome predictive modeling (CPM) (Shen et al. 2017) is a predictive modeling method that has been used widely in fMRI with a variety of outcome variables (Finn et al. 2015; Rosenberg et al. 2016; Yoo et al. 2018; Beaty et al. 2018 Jan 10; Lake et al. 2018 Mar 28). In brief, CPM first correlates each edge in the connectome with the phenotype of interest to identify edges that are associated with the outcome above some prespecified level (e.g., Pearson’s correlation with significance of *p* < 0.01). Out of these identified edges, the positively and negatively correlated edges are separately summed. These two values serve as inputs for two separate linear models (CPM+, CPM-) to predict the outcome scores. Of note, more recent studies have proposed a large number of other variants of CPM that use other methods of feature selection and feature aggregation, e.g., (Greene et al. 2018; Gao et al. 2019). We did not evaluate these methods in this study (see Limitations below).

#### Brain Basis Set (BBS)

Brain Basis Set (BBS) is a predictive modeling approach developed and validated in our previous studies (Sripada, Angstadt, et al. 2019; Kessler et al. 2016; Sripada, Rutherford, et al. 2019; Sripada, Angstadt, et al. 2020); see also studies by Wager and colleagues for a broadly similar approach (Wager et al. 2013; Chang et al. 2015; Woo, Schmidt, et al. 2017). BBS is similar to principal component regression (Park 1981; Jolliffe 1982), with an added predictive element. In a training partition, PCA is performed on an *n* subjects by *p* connectivity features matrix using the PCA function from scikit-learn in Python, yielding components ordered by descending eigenvalues. Expression scores are then calculated for each of *k* components for each subject by projecting each subject’s connectivity matrix onto each component. A linear regression model is then fit with these expression scores as predictors and the phenotype of interest as the outcome, saving **B**, the *k x 1* vector of fitted coefficients, for later use. In a test partition, the expression scores for each of the *k* components for each subject are again calculated. The predicted phenotype for each test subject is the dot product of **B** learned from the training partition with the vector of component expression scores for that subject. We tested two regimes for choosing the parameter k. In the first, k was set at 75 based on prior studies (Sripada, Angstadt, et al. 2019) which used dimensionality estimation algorithms to estimate the actual dimensionality of the HCP dataset and which showed that larger values tend to result in overfitting and worse performance. In the second regime, k was selected on each fold of our 10-fold cross validation (see below), using 5-fold cross validation within the training data.

#### Support Vector Regression (SVR)

SVR (Smola and Schölkopf 2004) finds a function *f* under the criteria that every data point fall within a required accuracy (*ε*) from *f* and *f* be as flat as possible. For linear functions *f*, this can be described as a convex optimization problem:

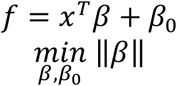

subject to:

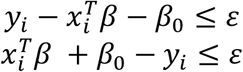

In cases where this optimization problem is infeasible, slack is added to the required accuracy. Support Vector regression allows *f* to be linear or non-linear in the feature space. The required accuracy is a model hyperparameter, and there is an additional hyperparameter C which further regulates the smoothness of *f*. Both hyperparameters were selected via a grid search on each fold of our 10-fold cross validation (see below), using 5-fold cross validation within the training data.

#### Random Forest

Random Forest regression (Liaw and Wiener 2002) averages the predictions from a collection of tree-based predictive models. Each tree in the collection is constructed using a different bootstrapped sample of the data and each split-point in the tree is chosen to be the most predictive feature from a random subset of all features. The predictions generated by tree-based models are noisy and susceptible to overfitting, and thus Random Forest regression benefits from averaging over the collection of these predictions. The maximum allowed depth of each tree in the collection and the number of features accessible at split-points are hyperparameters of the Random Forest model. These hyperparameters were selected via a grid search on each fold of our 10-fold cross validation (see below), using 5-fold cross validation within the training data.

### 2.7. HCP Outcome Variables

We used a total of 61 outcome variables from the HCP dataset (choice of these variables was guided by Kong et al. (2018), and a list of these variables is available in the Supplement). Two outcome variables were derived from factor analysis of HCP variables, and they are discussed in detail in our previous report (Sripada, Angstadt, et al. 2019). In brief, a general executive factor was created based on overall accuracy for three tasks: *n*-back working memory fMRI task, relational processing fMRI task, and Penn Progressive Matrices task. A speed of processing factor was created based on three NIH toolbox tasks: processing speed, flanker task, and card sort task, similar to a previous report (Carlozzi et al. 2015).

### 2.8 Train/Test Split and Calculation of ICC for Predictive Models

All predictive models were trained and tested in a 10-fold cross validation scheme. We calculated ICCs for predicted outcomes for these predictive models as follows (Figure 1): On each fold, we trained a predictive model on the train partition for session 1 data. We then used this trained model to generate predicted outcomes for the test partition in both session 1 and session 2, and we calculated the ICC of this pair of predicted outcomes. We next did this same procedure in the other direction: We trained the predictive model on the train partition for session 2 data, generated predicted outcomes for the test partition in both session 2 and session 1, and calculated their ICC. We repeated this process on all 10 folds and averaged over all 20 ICCs (2 for each fold).

**Figure 1:**
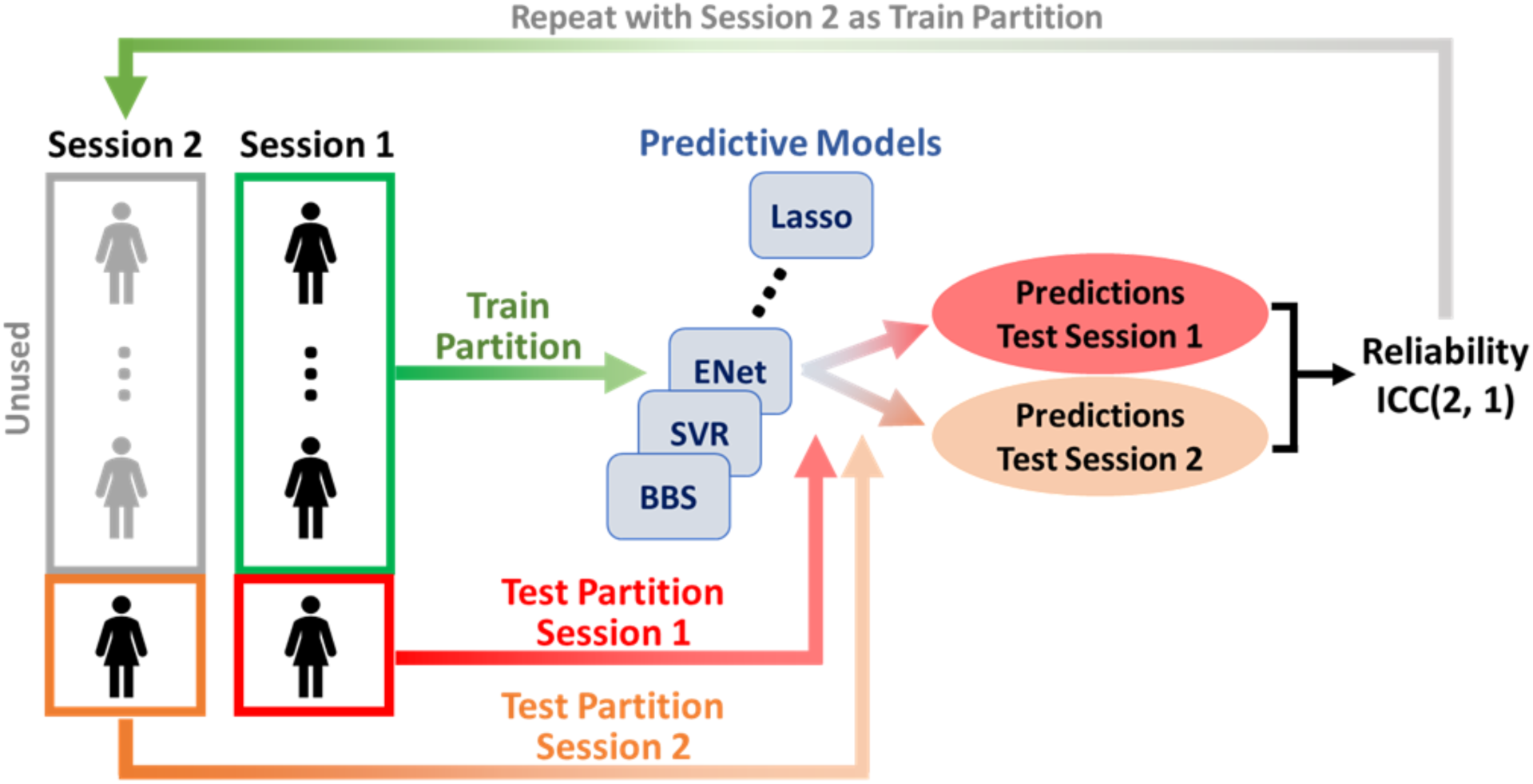
Overall Pipeline for Calculating Test-Retest Reliability of Predicted Outcomes of Predictive Models.

For the predictive modeling methods that have an additional tunable parameter (see the descriptions above), we tuned this parameter in an embedded cross-validation procedure within the train partition. But otherwise, we followed the procedure described above.

## 3 Results

### 3.1 For All Ten Predictive Modeling Methods, Mean Reliability of Predicted Outcomes Across 61 Phenotypes Was Higher Than Mean Reliability for Individual Connections

Figure 1 shows the test-retest reliabilities using functional connectomes generated from 29-minutes of data with 0.5mm FD motion scrubbing. The left side of Figure 1 shows the reliabilities for the individual connections of grayordinate (lighter shade) and volumetric connectomes (darker shade). Mean reliability (ICC) across individual edges was 0.50 for volumetric data and 0.35 for grayordinate data, but the spread was remarkably wide (standard deviation 0.19, 0.20, respectively). The next ten pairs of plots show test-retest reliabilities for the predicted outcomes of predictive models trained on 61 HCP outcome variables for grayordinate (lighter shade) and volumetric (darker shade) data. Grayordinate data consistently had lower test-retest reliability, which might be attributable to higher native resolution of surface-processed data or to the different parcellations applied to each (Power parcellation for volume and Gordon parcellation for grayordinate). Of note, all of the predictive modeling methods had higher mean test-retest reliability than individual edges.

Shou et al. proposed a whole-brain multivariate intraclass correlation coefficient for imaging maps, I2C2 (Shou et al. 2013). In short, I2C2 generalizes the univariate ICC to the case where the measurand is an image. The I2C2 reliability estimates for the volumetric and grayordinate whole brain connectomes (i.e., the connectomes used in producing the results in Figure 1) were 0.64 and 0.51, respectively. Seven (for volume) and all (for grayordinate) of the ten predictive models had higher mean test-retest reliability of predicted outcomes than the whole-connectome I2C2 coefficient.

Tables 1 and 2 show summary statistics for volumetric and grayordinate test-retest reliability as well as predictive accuracy for the ten predictive modeling methods. We in addition divided the 61 outcome variables into two categories: 1) Cognitive variables, which were derived from tasks from NIH Toolbox and Penn Neurocognitive Battery, as well as fMRI working memory task (N-back) and fMRI math calculation (math condition of the Language Task); and 2) All the other (non-cognitive) variables. For all ten predictive modeling methods, predictive accuracy was notably higher for cognitive variables.

**Table 1:**
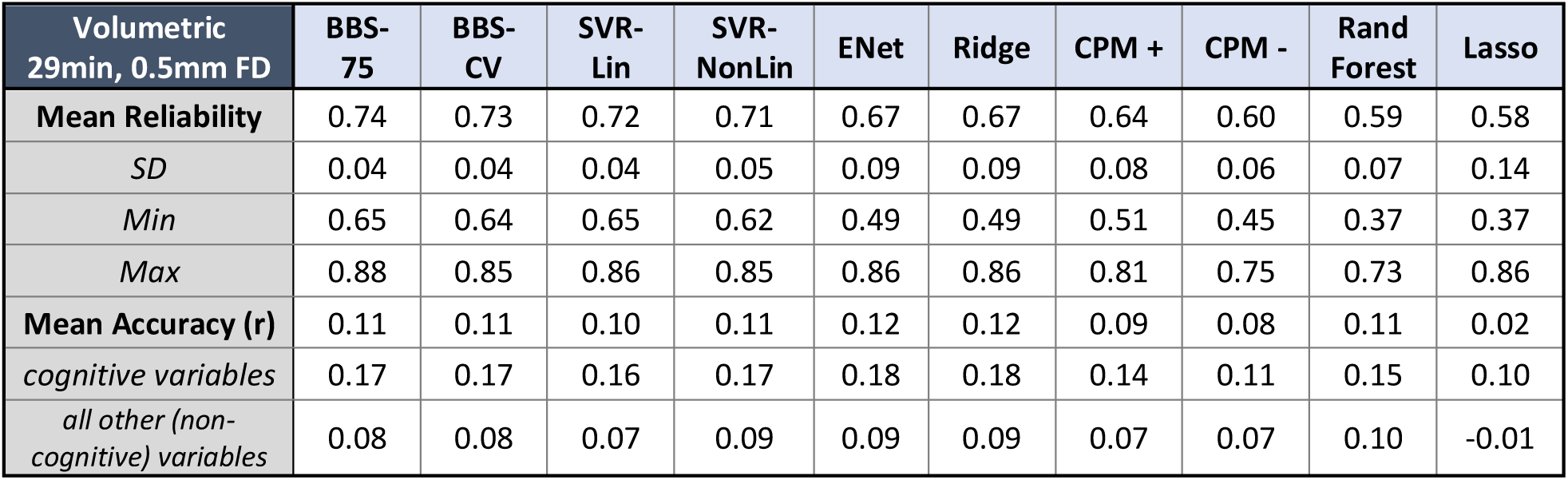
Summary Statistics for Test-Retest Reliability and Accuracy of Predictive Models Trained on Volumetric Data. Reliability is measured with the intraclass correlation (ICC) statistic. Accuracy is measured with Pearson’s correlation between each predicted and actual outcome variable. Interestingly cognitive variables, mostly from the NIH Toolbox and Penn Neurocognitive Battery, consistently yielded better accuracy than all other (non-cognitive) variables.

| Volumetric<br>29min, 0.5mm FD | BBS-75 | BBS-CV | SVR-Lin | SVR-NonLin | ENet | Ridge | CPM + | CPM - | Rand<br>Forest | Lasso |
| --- | --- | --- | --- | --- | --- | --- | --- | --- | --- | --- |
| Mean Reliability | 0.74 | 0.73 | 0.72 | 0.71 | 0.67 | 0.67 | 0.64 | 0.60 | 0.59 | 0.58 |
| SD | 0.04 | 0.04 | 0.04 | 0.05 | 0.09 | 0.09 | 0.08 | 0.06 | 0.07 | 0.14 |
| Min | 0.65 | 0.64 | 0.65 | 0.62 | 0.49 | 0.49 | 0.51 | 0.45 | 0.37 | 0.37 |
| Max | 0.88 | 0.85 | 0.86 | 0.85 | 0.86 | 0.86 | 0.81 | 0.75 | 0.73 | 0.86 |
| Mean Accuracy (r) | 0.11 | 0.11 | 0.10 | 0.11 | 0.12 | 0.12 | 0.09 | 0.08 | 0.11 | 0.02 |
| cognitive variables | 0.17 | 0.17 | 0.16 | 0.17 | 0.18 | 0.18 | 0.14 | 0.11 | 0.15 | 0.10 |
| all other (non-cognitive) variables | 0.08 | 0.08 | 0.07 | 0.09 | 0.09 | 0.09 | 0.07 | 0.07 | 0.10 | -0.01 |

**Table 2:** Summary Statistics for Test-Retest Reliability and Accuracy of Predictive Models Trained on Grayordinate Data. Comparing volumetric and grayordinate results, all ten predictive models had greater mean reliability of predicted outcomes when trained on volumetric data. In contrast, all predictive models except Random Forest showed greater mean accuracy when trained using grayordinate data.

| Grayordinate<br>29min, 0.5mm FD | BBS-75 | BBS-CV | SVR-Lin | SVR-NonLin | ENet | Ridge | CPM + | Lasso | CPM - | Rand<br>Forest |
| --- | --- | --- | --- | --- | --- | --- | --- | --- | --- | --- |
| Mean Reliability | 0.65 | 0.65 | 0.64 | 0.63 | 0.62 | 0.62 | 0.58 | 0.58 | 0.55 | 0.53 |
| SD | 0.05 | 0.05 | 0.04 | 0.06 | 0.07 | 0.07 | 0.07 | 0.12 | 0.05 | 0.08 |
| Min | 0.56 | 0.53 | 0.54 | 0.54 | 0.48 | 0.46 | 0.45 | 0.30 | 0.46 | 0.30 |
| Max | 0.82 | 0.81 | 0.82 | 0.82 | 0.83 | 0.83 | 0.75 | 0.80 | 0.68 | 0.72 |
| Mean Accuracy (r) | 0.13 | 0.13 | 0.12 | 0.13 | 0.13 | 0.13 | 0.10 | 0.07 | 0.09 | 0.11 |
| cognitive variables | 0.20 | 0.19 | 0.18 | 0.18 | 0.20 | 0.20 | 0.14 | 0.13 | 0.11 | 0.14 |
| all other (non-cognitive) variables | 0.11 | 0.10 | 0.10 | 0.11 | 0.11 | 0.11 | 0.08 | 0.04 | 0.08 | 0.09 |

Tables 1 and 2 show that several predictive modeling methods had consistently high levels of performance across the 61 phenotypes. Reliability is conventionally classified as follows: 0– 0.4 = poor, 0.4–0.6 = fair, 0.6–0.75 = good and 0.75–1 = excellent (Cicchetti 1994; Cicchetti and Sparrow 1981). For the 29-minutes of volumetric data with 0.5mm FD thresholding, several predictive modeling methods exceeded the 0.60 threshold for all phenotypes, namely: BBS-75, BBS-CV, SVR-Lin, and SVR-NonLin. Other models, including Ridge, Elastic Net, CPM-Positive, Random Forest, Lasso and CPM-Negative, exceeded this threshold more inconsistently (48, 48, 39, 29, 25, and 24 out of 61 phenotypes respectively). For the 29-minutes of grayordinate data, good but consistently weaker reliability performance was observed. The strongest performing models here, BBS-75, BBS-CV and SVR-Linear, exceeded the threshold for ‘good’ reliability for 54, 52, and 54 out of 61 phenotypes, respectively.

### 3.2 Reliability of Predicted Outcomes Shows Sensitivity to Scan Length, but is Unaffected by Motion Scrubbing Threshold

Several studies have reported associations between scan-length and the reliability of functional connectivity (Birn et al. 2013; Elliott et al. 2020). To evaluate the sensitivity of test-retest reliability for predicted outcomes of predictive models to scan-length, we employed functional connectivity matrices generated using 29-, 15- and 7.5-minutes of run data from each day. In addition, recent studies have also highlighted the effects of motion-related artifacts on resting-state connectivity (Power et al. 2012; Satterthwaite et al. 2012; Van Dijk et al. 2012). Thus, we also considered two framewise displacement thresholds for censoring frames: 0.5mm and 0.2mm.

Results (Figure 3) showed that for volumetric data with 0.5mm FD thresholding, the number of predictive models with mean reliability greater than 0.60 drops from seven out of ten for 29- and 15-minutes of data to just three out of ten for 7.5-minutes of data. Similarly, for grayordinate data with 0.5mm FD thresholding, the number of models exceeding 0.60 mean reliability is six, one and zero out of ten for 29-, 15-, and 7.5-minutes, respectively. In contrast, however, more stringent motion scrubbing shows minimal effect on test-retest reliability of predictive models, suggesting motion-related artifacts are not key contributors to predictive model reliability. Finally, this pattern of results for test-retest reliability (i.e., sensitivity to scan length but not motion scrubbing threshold) is broadly mirrored in results for accuracy of predictive models (Figure 4).

**Figure 2:**
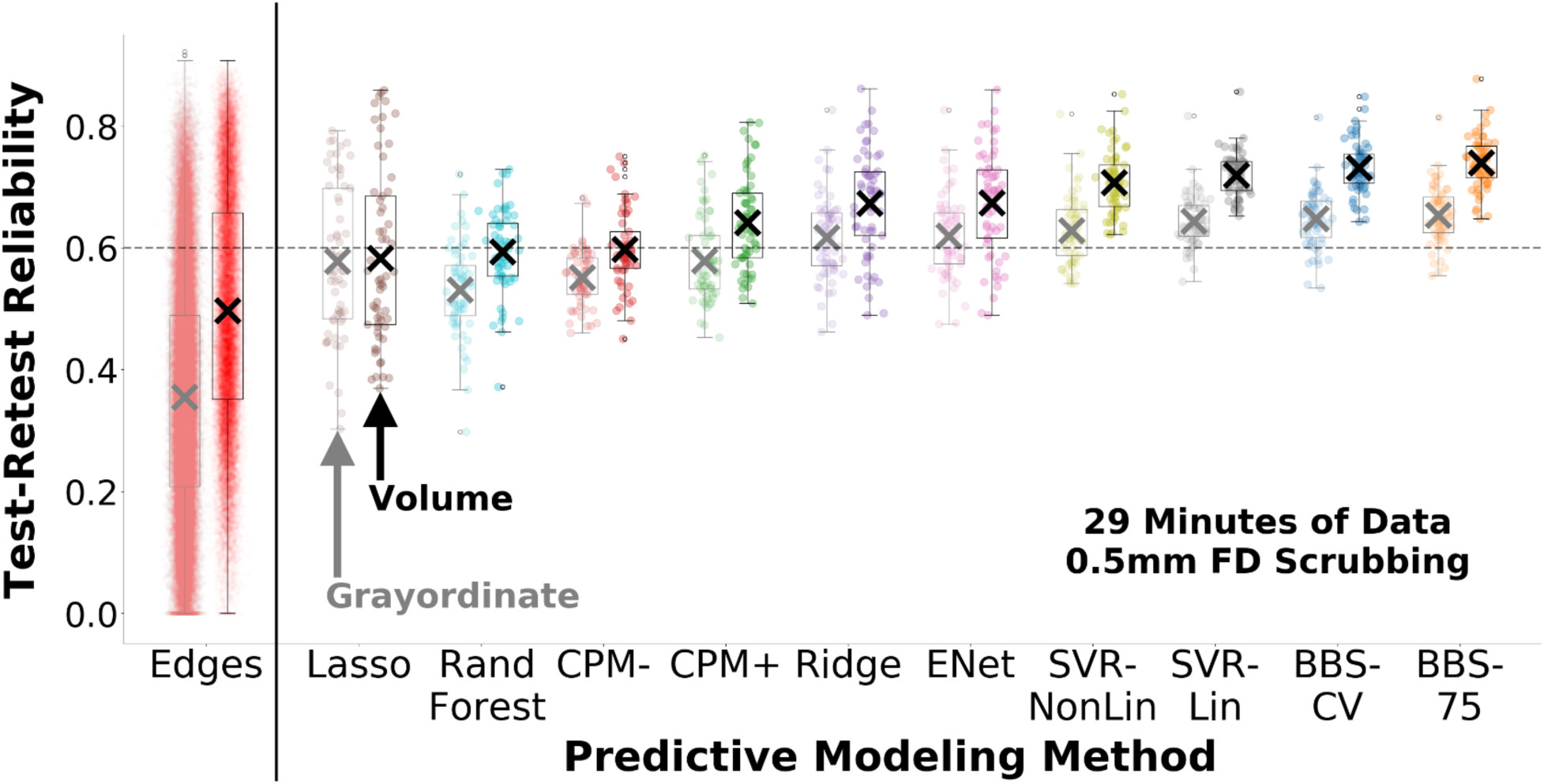
Distribution of Test-Retest Reliabilities for Individual Connections and for Predicted Outcomes of Predictive Models for Volumetric and Grayordinate Data. Reliabilities for individual connections were calculated for all connections in the resting state connectome. Mean reliability for individual connections was relatively low, but the range was wide. Reliabilities for predicted outcomes of predictive models were calculated for 61 different outcome variables available in the HCP dataset. Seven (for volume) and six (for grayordinate) of ten predictive modeling methods had mean reliabilities of predicted outcomes greater than 0.60 (marked by the dashed line), conventionally classified as good.

**Figure 3:**
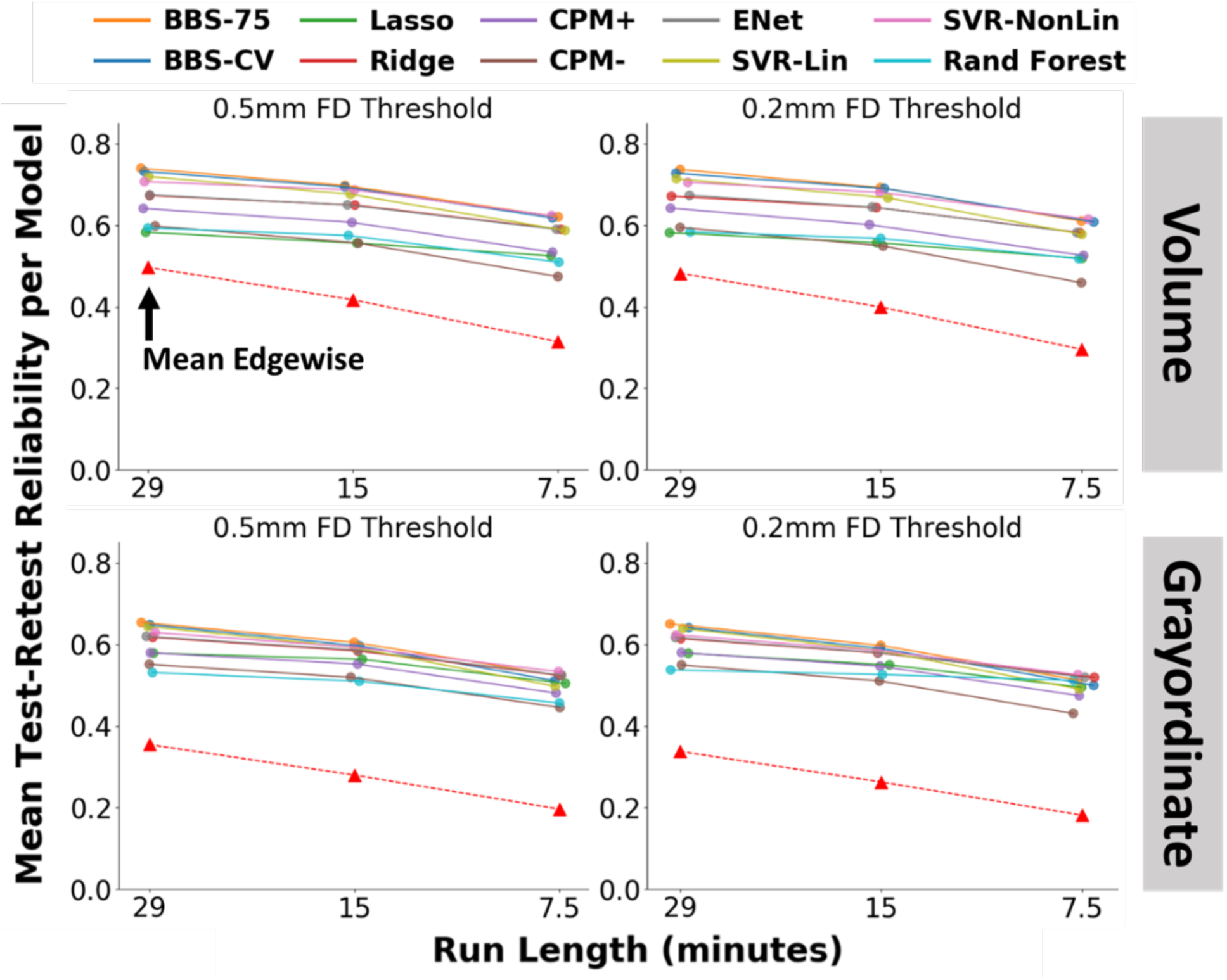
Test-Retest Reliabilities for Predicted Outcomes of Predictive Models Across Run-Lengths and FD Thresholds. The mean test-retest reliability of predictive models across the 61 phenotypes was computed for three different run-lengths and two motion scrubbing thresholds, for volume and grayordinate data. Reliability is highest at the longest scan duration (29 mins) and drops consistently with shorter scan durations, but it is minimally affected by extent of motion scrubbing. A sizable boost in test-retest reliability (compared to reliability of individual edges of the connectome; red triangles in the figure) was observed predictive models across all processing methods, scan lengths, and motion scrubbing thresholds.

**Figure 4:**
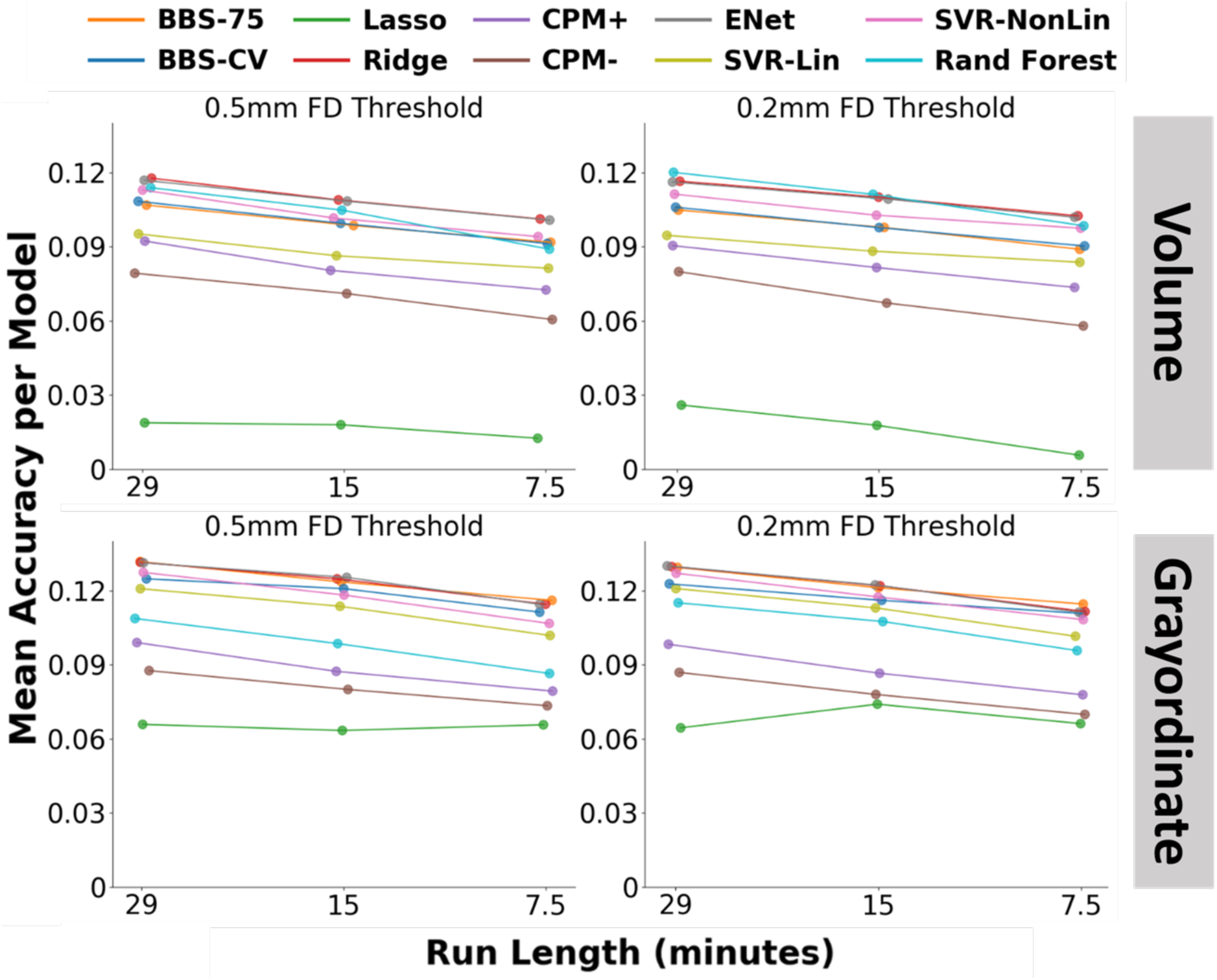
Accuracies of Predicted Outcomes of Predictive Models Across Run-Lengths and FD Thresholds. The mean accuracy of predictive models across the 61 phenotypes was computed for three different run-lengths and two motion scrubbing thresholds, for volume and grayordinate data. Accuracy is highest at the longest scan duration (29 mins) and drops consistently with shorter scan durations, but it is minimally affected by extent of motion scrubbing.

### 3.3. Multiple Mechanisms Likely Explain Why Predictive Models Have a Large Boost in Test-Retest Reliability

We next examined several mechanisms that could help to explain why predicted outcomes of predictive models are more reliable than individual features.

#### Selection of High Variance Features

Test-retest reliability is positively related to inter-subject variance of features (Nunnally 1970; Woo and Wager 2016). BBS uses dimensionality reduction with PCA, an algorithm that finds components with maximal variance explained (Abdi and Williams 2010). Thus, one mechanism that could improve reliability, a mechanism that is specific to BBS (e.g., BBS-75 and BBS-CV), is use of features with high inter-subject variance. Consistent with this idea, we found that with 29-minutes of volumetric data (0.5 FD threshold), the mean variance of the individual connections in the connectome was 0.01. But mean variance of expression scores of the first 75 PCA components is 1.60, more than 150 times larger.

#### Selection of Correlated Features

Another mechanism that could improve reliability for nearly all the predictive modeling methods we tested is selecting correlated features and aggregating over them. It is well known from classical test theory that the sum of a set of positively correlated features will have higher reliability than the features themselves (Nunnally 1970) (see the Supplement for a general equation linking reliability of a weighted sum to the statistical properties of the individual features). In our previous work (Sripada, Angstadt, et al. 2019), we used multiple convergent methods to establish that there are substantial inter-correlations in connectivity features that differ across individuals. Because of this high dependence, just 50 to 150 components capture most meaningful individual differences. BBS specifically selects correlated features through the use of PCA: Each component consists of a weighted set of features that are jointly co-expressed across subjects (in proportion to the component expression), and thus these features are correlated across subjects (Abdi and Williams 2010). Supporting this idea, we found that with 29-minutes of volumetric data (0.5 FD threshold), the mean test-retest reliability of expression scores of the first 75 PCA components is 0.72. With this same data, mean reliability of individual features is 0.50, considerably lower. Of note, this boost in reliability for components likely reflects both the operation of this second mechanism (selecting correlated features) as well as the first (selecting high variance features).

Several of the other predictive modeling methods also select correlated features, but in different ways. For example, CPM performs a search for features that are correlated with the outcome variable up to a desired level of statistical significance (e.g., p< 0.01). Since these features are all correlated with the behavioral variable of interest, they will also tend to be correlated with each other as well. Consistent with this idea, we found that with 29-minutes of volumetric data (0.5 FD threshold), mean pairwise intercorrelation of all features across the connectome is 0.015, while the mean pairwise intercorrelation of CPM-selected feature set is 0.071 for CPM positive and 0.079 for CPM negative.

Ridge regression selects correlated features as well. The operation of Ridge Regression is best understood in contrast with Lasso. When several collinear predictors all predict an outcome well, Lasso’s L_1_ penalty favors sparsity and tends to shrink the coefficients of all but one of the predictors to zero. Ridge Regression’s L_2_ penalty, in contrast, will tend to assign large and highly similar regression weights to these colinear features. In this way, Ridge Regression “selects” (or more correctly, weights more heavily) these highly intercorrelated features. Of note, Elastic Net attempts to balance the L_1_ and L_2_ penalties. Nonetheless, in practice it often strongly favors the ridge penalty, as was the case in this study: With 29-minutes of volumetric data (0.5 FD threshold), the mean Pearson’s correlation between the predicted outcomes from Elastic Net and Ridge Regression were 0.996 for volumetric and 0.993 for grayordinate (standard deviations = 0.030, 0.049 respectively).

#### Selection of Valid Features

Let us make the following assumption: There in fact is a sizable quantity of true variance in the connectome, i.e., variance that relates to an outcome variable of interest. While some of this true variance may be state specific, it is plausible that some of it will not be; instead it will reflect stable, non-noise connectomic differences that are related to the outcome variable. All of the predictive modeling methods we examined select features that are correlated with a specified outcome variable. Given our assumption, then, features selected by these predictive modeling methods will correspondingly be enriched with respect to these valid, stable connectomic differences. This enrichment with respect to true variance will boost test-retest reliability.

To further assess the role of this third mechanism in boosting reliability, we examined the relationship for each predictive modeling method between accuracy in predicting a phenotype (which is expected to reflect quantity of valid variance in the phenotype) and observed reliability of predicted scores for that phenotype. A positive relationship would suggest that when a phenotype contains more valid variance (indexed by accuracy), predicted scores from predictive models for the phenotype will exhibit higher test-retest reliability. We did in fact observe large positive relationships for nine of ten predictive modeling methods (all except CPM-) that ranged from *r*=0.40 for SVR-Lin to *r*=0.83 for Lasso (Figure 5). This result is consistent with the idea that predictive modeling methods identify and leverage valid variance in the connectome, and this plays a role in boosting reliabilities for predicted scores. Of note, interpretation of the preceding finding requires some caution, since the observed correlation could also be explained in the reverse direction: to the extent predictive models leverage more reliable features, their predicted outcomes are more reliable, and this in turn tends to improve their predictive accuracy.

**Figure 5:**
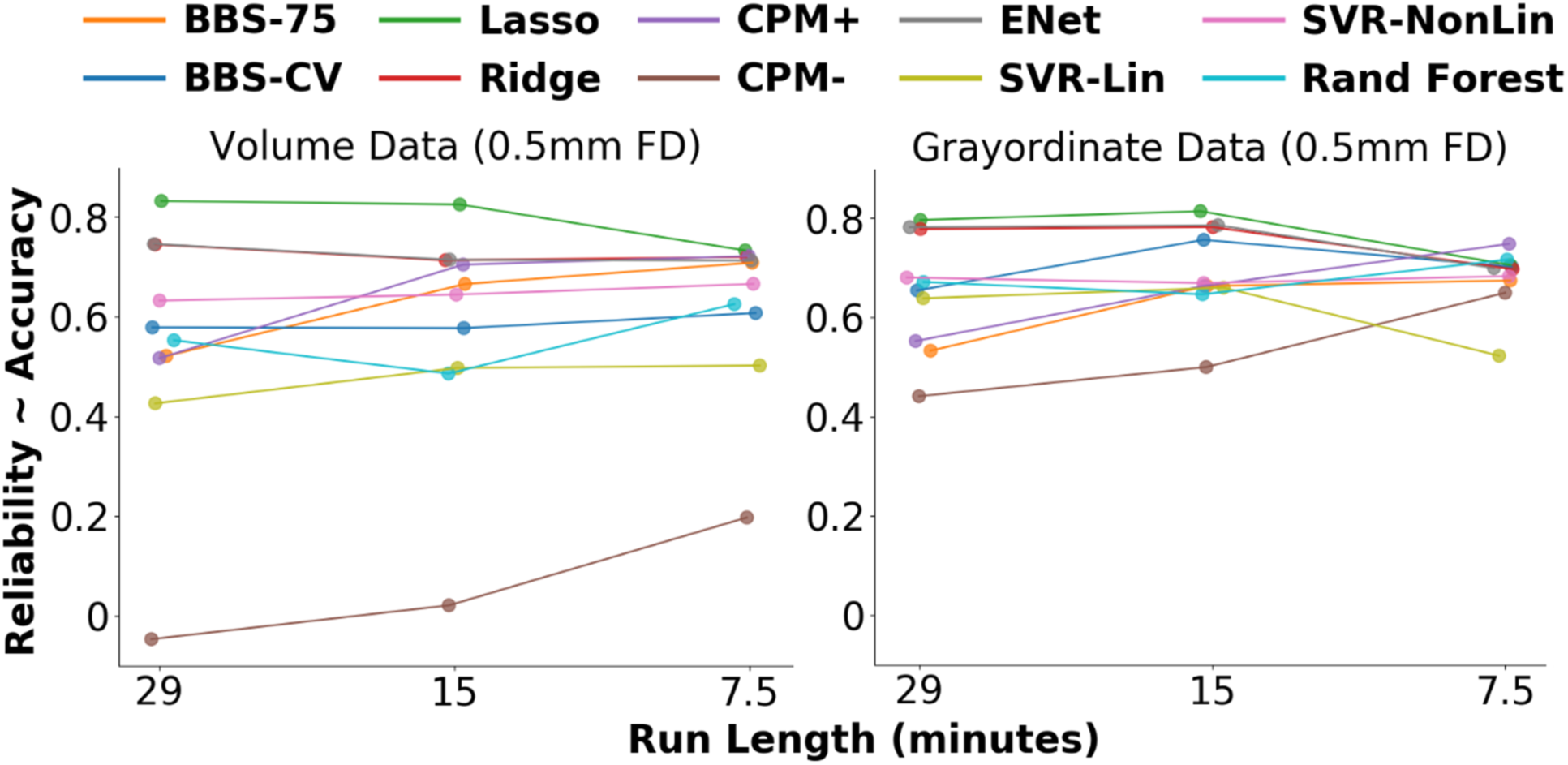
Relationship Between Test-Retest Reliability and Accuracy of Predicted Outcomes of Predictive Models. We hypothesized that the presence of more valid variance in the connectome for a phenotype would improve test-retest reliability, leading to a positive relationship between accuracy of predicted outcomes of predictive models and their test-retest reliability. Thus, we calculated the correlation between accuracy (measured with Pearson’s r between predicted versus actual outcomes) and reliability across 61 HCP phenotypes. For nine out of ten predictive modeling methods, we observed large positive correlations.

## 4 Discussion

This is the first study to systematically investigate test-retest reliability of multivariate predictive models applied to resting state connectivity maps. We found that in contrast to reliability of individual resting state connections, reliability of the predicted outcomes of predictive models is substantially higher for all ten predictive modeling methods that we examined. Moreover, this sizable improvement was observed across all processing methods, scan lengths, and motion censoring thresholds. For the most reliable methods, including Brain Basis Set and Support Vector Regression, the mean reliability of predicted outcomes was consistently in the “good” range (above 0.60) across nearly all scanning and processing parameters. Test-retest reliability is critical for the use of fMRI in individual differences research. Our results suggest more widespread use of predictive models can help address questions about reliability that have been raised in recent reports and that remain a serious concern for the neuroimaging field.

### Test-Retest Reliability and Units of Analysis

Previous studies of reliability in resting state fMRI have mostly examined individual connections (Shehzad et al. 2009; Birn et al. 2013; Noble et al. 2017). While results have varied, a recent meta-analysis (Noble et al. 2019) found reliability was typically relatively low at 0.29. Broadly consistent with this result, we found mean reliability of individual connections in the HCP dataset ranged from 0.18 to 0.50, depending on scanning and processing parameters (see Figure 3). Several studies examined larger, more complex units of analysis and found higher levels of reliability. For example, Noble and colleagues (Noble et al. 2017) examined mean connectivity within intrinsic connectivity networks such as default mode network and fronto-parietal network. They found reliabilities were modestly higher for networks than for individual connections (range 0.35 to 0.60 for networks). Similarly, a modest boost in reliability appears to be observed with higher-order metrics such as graph theory metrics (Braun et al. 2012; Termenon et al. 2016). Predictive models, which aggregate across a still wider range of features to produce whole-connectome neurosignatures, arguably represent a still larger, more complex unit of analysis. In the present study, we found clear evidence of substantially higher test-retest reliability for predicted outcomes of predictive models for all ten methods tested. Overall, these results suggest that test-retest reliability differs substantially across both size of the units of analysis (i.e., edge-, network-, neurosignature-level), as well as the types of aggregation methods that were utilized to generate these higher-level units.

### Should We Be Pessimistic or Optimistic About Using Resting State Connectivity for Individual Differences Research?

Given high mean reliability for predicted outcomes of predictive models and much lower mean reliability for individual connections, should we be optimistic or pessimistic about reliability of resting state connectivity? While we acknowledge both perspectives capture part of the overall picture, we briefly suggest there is more reason for optimism.

Test-retest reliability is most critical for research that seeks to use imaging features to predict individual differences, for example, translational neuroimaging research that aims to construct brain-based biomarkers. This is because reliability is mathematically related to predictive validity, the ability of measures to predict an outcome. According to an important formula in classical test theory, reliability sets a ceiling on predictive validity, and as reliability falls, the maximum achievable predictive validity drops with it (Nunnally 1970).

But, critically, if one’s goal is in fact prediction of individual differences of some outcome variable of interest (e.g., behaviors or symptoms), focusing on individual connections of the connectome is unlikely to be a fruitful approach. This is because for most constructs of interest to translational researchers (e.g., general cognitive ability, neuroticism, pain, autism), it is unlikely that any single connection contains much discriminative information about the construct. Rather, it is likely that this discriminative information resides in distributed changes across widespread network connections (Dubois et al. 2018; Sripada, Rutherford, et al. 2019; Sripada, Angstad, et al. 2020). It is thus plausible that multivariate predictive models, which aggregate information from across the connectome, will produce larger effect sizes than univariate methods (Reddan et al. 2017; Woo, Chang, et al. 2017). Indeed, in studies that examined both head-to-head in resting state fMRI (Marek et al. 2020) and task-based fMRI (Chang et al. 2015; Woo and Wager 2016; Woo, Schmidt, et al. 2017), predictive models outperformed individual imaging features.

In short, then, predictive models are arguably a more important tool for individual differences research in fMRI than univariate tests applied to individual imaging features. If this is correct, then poor reliability of individual imaging features may not be a decisive concern. Rather, a more optimistic interpretation is available: Predictive models are the critically important tools we need for individual differences research in neuroimaging, and at least some predictive modeling methods *do* appear to have adequate levels of test-retest reliability.

### Why Predictive Models Have Better Test-Retest Reliability

We identified three factors that can explain why predictive models have better reliability compared to individual connections: selection of high variance features, correlated features, and valid features. Notably, these factors reflect basic statistical properties of predictive modeling methods and thus would be expected to operate with great generality. Consistent with this idea, we observed a sizable boost in test-retest reliability for predicted outcomes of predictive models across a wide variety of phenotypes (cognition, emotion, personality, and psychopathology) and across a wide variety of data types (volumetric versus surface-based processing, different scan lengths, and different motion censoring thresholds). Notably, the size of the boost conferred by predictive models tended to be largest at the shortest scanning sessions with surface-based data, which is where edge-level reliability is at its lowest. That is, predictive models appear to help reliability the most precisely where help is most needed.

### Implications for Task fMRI

Recent reports also find poor reliability in task fMRI (Elliott et al. 2020). This result may be seen as particularly discouraging because many researchers have thought that tasks, because they involve carefully controlled manipulations of psychological constructs, might be an especially effective way of detecting differences in these constructs across individuals (Matthews et al. 2006). Could predictive models play a similar role in boosting reliability with task-based fMRI? As noted above, the three mechanisms we identified for why predictive models boost reliability are quite general and reflect basic statistical properties of predictive modeling methods, suggesting that these mechanisms would indeed also be operative with task activation maps. Moreover, one study found initial evidence that predictive models do in fact produce a boost in reliability in the task setting: Woo and Wager (2016) report that model-based predictions of pain ratings during a nociceptive stimulation task had higher reliability (ICC=0.72) than three regions of interest known to be associated with pain (ICCs: 0.54 to 0.59). Future studies should extend the present work to systematically assess reliability of predicted outcomes of predictive models applied to task data.

An additional connection between the present study and task fMRI concerns the use of task fMRI to generate functional connectivity matrices, and there are at least two importantly different approaches. First, task-related signals could be regressed out of task time series data, e.g., (Fair et al. 2007), to generate additional “resting state-like” data. Given our finding that predictive models have higher test-retest reliability at longer scan durations, utilizing task data in this way would be expected to further boost reliability. Second, task fMRI data could be used to generate task-based functional connectivity matrices. There is some evidence that these task connectomes have better test-retest reliability and improved predictivity for phenotypes of interest (Greene et al. 2018; Elliott et al. 2019). Thus, applying predictive modeling methods to task-based connectomes could provide an additional avenue to further boost test-retest reliability.

### Limitations

This study has several limitations. First, we assessed ten popular predictive modeling methods. Nonetheless, there are a large number of other predictive modeling methods that we did not study, and future work should systematically compare them. Second, while we examined a large number of outcome variables (61 in total), there are of course a vast number of outcome variables that we could not test with the HCP dataset (e.g., pain ratings, schizophrenia status, depression-treatment response, etc.). As more comprehensive datasets become available, it would be useful to extend these results to a still broader range of outcome variables. It would additionally be useful to study datasets with longer temporal separation between test and retest sessions. Third, our analyses in this study essentially assume that 61 phenotypic measures that served as outcome variables are measured with perfect reliability. In reality, these variables are measured with varying degrees of noise, and interpretation of the reliability of predictive models should account for the fact that their reliability will be reduced because the predictive target is itself noisy. Fourth, it bears emphasis that test-retest reliability is a statistic that is specific to a given population. Most relevant for the present purposes, it is highly sensitive to the inter-individual variance in imaging features (Nunnally 1970). The HCP dataset consists of a fairly homogenous sample of psychologically healthy young adults. It is possible that reliability will be higher in fMRI, at both the individual feature-level as well as the predictive model-level, if more heterogenous samples are considered, as this could potentially boost inter-individual variance in imaging features (Woo and Wager 2016). Finally, our focus in this manuscript has been on test-retest reliability of predicted outcomes of predictive models. A distinct but related question concerns the consistency of the weighting function over features learned by predictive models across two occasions of testing (temporal stability) or across two different groups of subjects (reproducibility). Answers to these questions are important for researchers interested in interrogating and interpreting the neurosignatures learned by predictive models, and thus the consistency of this neurosignature should be studied in future work.

## Conclusion

In sum, this study is the first to systematically assess the test-retest reliability of predicted outcomes of predictive models applied to resting state connectivity maps. In contrast to the somewhat bleak conclusions of recent studies about reliability of individual imaging features, we found that predicted outcomes of predictive modeling methods have much improved test-retest reliability.

## Data Availability

All data and code used to produce the results in this manuscript are available online: https://github.com/SripadaLab/Predictive_Modeling_Reliability_HCP

## Supplement

Composite Reliability Formula (He 2009)

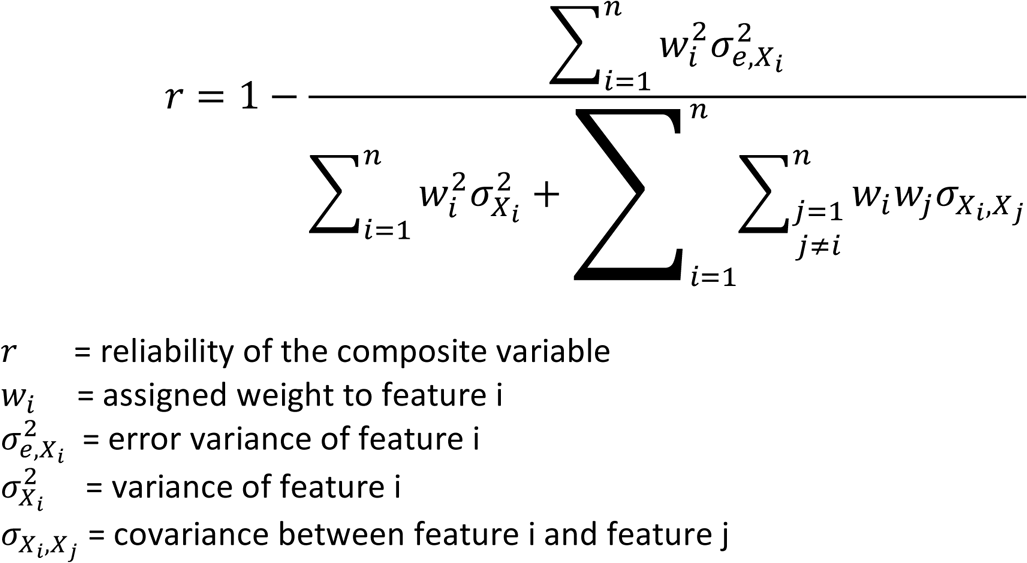

The composite reliability formula relates test-retest reliability of a composite variable to statistical properties of the variables that are summed. The formula highlights that test-retest reliability of a composite variable is positively related to the variables in the numerator, specifically: 1) the variance of the features that enter the composite; 2) the weights places on these features (i.e., larger weights placed on higher variance features yield higher reliability).

### Outcome Variables from HCP Dataset

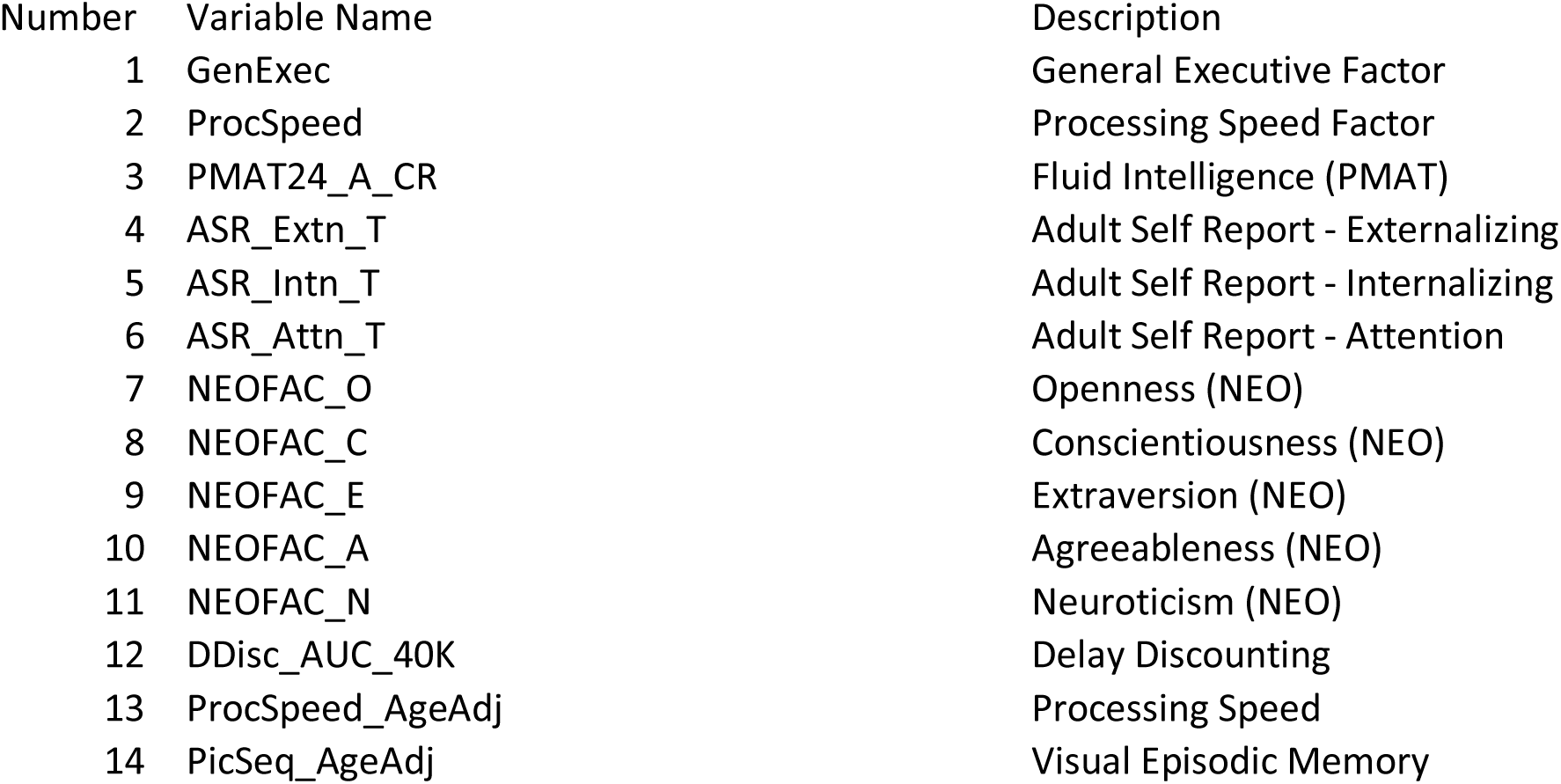
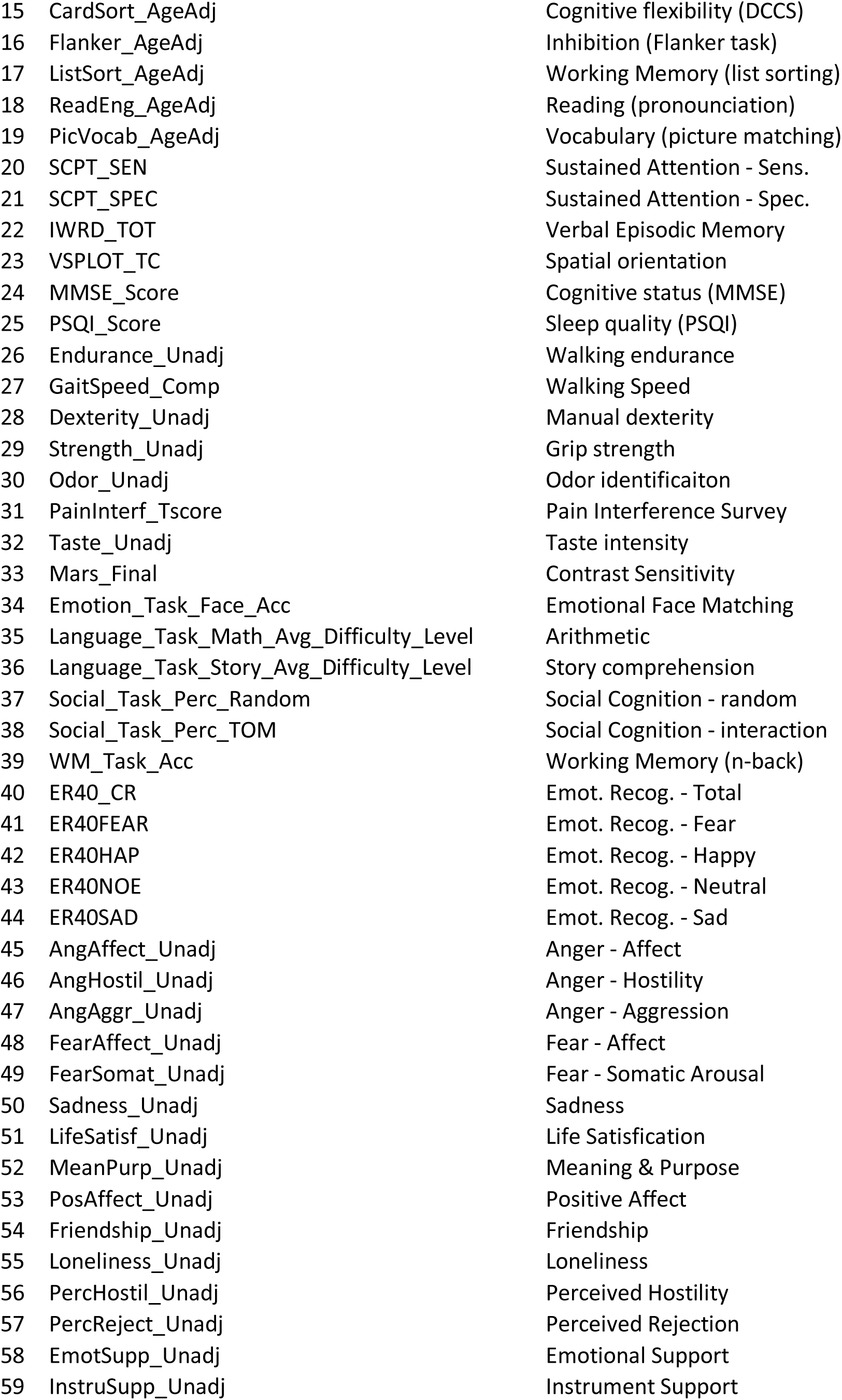
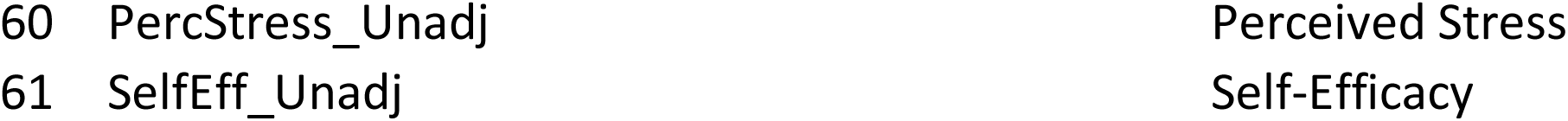

## References

Abdi H, Williams LJ. 2010. Principal component analysis. Wiley Interdiscip Rev Comput Stat. 2(4):433–459.

Beaty RE, Kenett YN, Christensen AP, Rosenberg MD, Benedek M, Chen Q, Fink A, Qiu J, Kwapil TR, Kane MJ, et al. 2018 Jan 10. Robust prediction of individual creative ability from brain functional connectivity. Proc Natl Acad Sci.:201713532. doi:10.1073/pnas.1713532115.

Behzadi Y, Restom K, Liau J, Liu TT. 2007. A component based noise correction method (CompCor) for BOLD and perfusion based fMRI. NeuroImage. 37(1):90–101. doi:10.1016/j.neuroimage.2007.04.042.

Birn RM, Molloy EK, Patriat R, Parker T, Meier TB, Kirk GR, Nair VA, Meyerand ME, Prabhakaran V. 2013. The effect of scan length on the reliability of resting-state fMRI connectivity estimates. NeuroImage. 83:550–558. doi:10.1016/j.neuroimage.2013.05.099.

Braun U, Plichta MM, Esslinger C, Sauer C, Haddad L, Grimm O, Mier D, Mohnke S, Heinz A, Erk S. 2012. Test–retest reliability of resting-state connectivity network characteristics using fMRI and graph theoretical measures. Neuroimage. 59(2):1404–1412.

Carlozzi NE, Beaumont JL, Tulsky DS, Gershon RC. 2015. The NIH Toolbox Pattern Comparison Processing Speed Test: Normative Data. Arch Clin Neuropsychol. 30(5):359–368. doi:10.1093/arclin/acv031.

Chang LJ, Gianaros PJ, Manuck SB, Krishnan A, Wager TD. 2015. A sensitive and specific neural signature for picture-induced negative affect. PLoS Biol. 13(6):e1002180.

Cicchetti DV. 1994. Guidelines, criteria, and rules of thumb for evaluating normed and standardized assessment instruments in psychology. Psychol Assess. 6(4):284.

Cicchetti DV, Sparrow SA. 1981. Developing criteria for establishing interrater reliability of specific items: applications to assessment of adaptive behavior. Am J Ment Defic.

Diedrichsen J, Maderwald S, Küper M, Thürling M, Rabe K, Gizewski ER, Ladd ME, Timmann D. 2011. Imaging the deep cerebellar nuclei: a probabilistic atlas and normalization procedure. Neuroimage. 54(3):1786–1794.

Drysdale AT, Grosenick L, Downar J, Dunlop K, Mansouri F, Meng Y, Fetcho RN, Zebley B, Oathes DJ, Etkin A. 2017. Resting-state connectivity biomarkers define neurophysiological subtypes of depression. Nat Med. 23(1):28.

Dubois J, Galdi P, Paul LK, Adolphs R. 2018. A distributed brain network predicts general intelligence from resting-state human neuroimaging data. Philos Trans R Soc B Biol Sci. 373(1756). doi:10.1098/rstb.2017.0284. http://rstb.royalsocietypublishing.org/content/373/1756/20170284.abstract.

Elliott ML, Knodt AR, Cooke M, Kim MJ, Melzer TR, Keenan R, Ireland D, Ramrakha S, Poulton R, Caspi A. 2019. General functional connectivity: Shared features of resting-state and task fMRI drive reliable and heritable individual differences in functional brain networks. NeuroImage.189:516–532.

Elliott ML, Knodt AR, Ireland D, Morris ML, Poulton R, Ramrakha S, Sison ML, Moffitt TE, Caspi A, Hariri AR. 2020. What Is the Test-Retest Reliability of Common Task-Functional MRI Measures? New Empirical Evidence and a Meta-Analysis. Psychol Sci.:0956797620916786.

Fair DA, Schlaggar BL, Cohen AL, Miezin FM, Dosenbach NUF, Wenger KK, Fox MD, Snyder AZ, Raichle ME, Petersen SE. 2007. A method for using blocked and event-related fMRI data to study “resting state” functional connectivity. NeuroImage. 35(1):396–405. doi:10.1016/j.neuroimage.2006.11.051.

Finn ES, Shen X, Scheinost D, Rosenberg MD, Huang J, Chun MM, Papademetris X, Constable RT. 2015. Functional connectome fingerprinting: identifying individuals using patterns of brain connectivity. Nat Neurosci. 18(11):1664–1671. doi:10.1038/nn.4135.

Gao S, Greene AS, Constable RT, Scheinost D. 2019. Combining multiple connectomes improves predictive modeling of phenotypic measures. Neuroimage. 201:116038.

Glasser MF, Sotiropoulos SN, Wilson JA, Coalson TS, Fischl B, Andersson JL, Xu J, Jbabdi S, Webster M, Polimeni JR, et al. 2013. The minimal preprocessing pipelines for the Human Connectome Project. NeuroImage. 80:105–124. doi:10.1016/j.neuroimage.2013.04.127.

Gordon EM, Laumann TO, Adeyemo B, Huckins JF, Kelley WM, Petersen SE. 2016. Generation and Evaluation of a Cortical Area Parcellation from Resting-State Correlations. Cereb Cortex N Y N 1991. 26(1):288–303. doi:10.1093/cercor/bhu239.

Greene AS, Gao S, Scheinost D, Constable RT. 2018. Task-induced brain state manipulation improves prediction of individual traits. Nat Commun. 9(1):1–13.

He Q. 2009. Estimating the reliability of composite scores. Coventry: Ofqual.

Hoerl AE, Kennard RW. 1970. Ridge regression: Biased estimation for nonorthogonal problems. Technometrics. 12(1):55–67.

Jolliffe IT. 1982. A note on the use of principal components in regression. Appl Stat.:300–303.

Kessler D, Angstadt M, Sripada C. 2016. Brain Network Growth Charting and the Identification of Attention Impairment in Youth. JAMA Psychiatry. 73(5):481–489.

Klöppel S, Abdulkadir A, Jack Jr CR, Koutsouleris N, Mourão-Miranda J, Vemuri P. 2012. Diagnostic neuroimaging across diseases. Neuroimage. 61(2):457–463.

Kong R, Li J, Orban C, Sabuncu MR, Liu H, Schaefer A, Sun N, Zuo X-N, Holmes AJ, Eickhoff SB. 2018. Spatial topography of individual-specific cortical networks predicts human cognition, personality, and emotion. Cereb Cortex. 29(6):2533–2551.

Lake EMR, Finn ES, Noble SM, Vanderwal T, Shen X, Rosenberg MD, Spann MN, Chun MM, Scheinost D, Constable RT. 2018 Mar 28. The functional brain organization of an individual predicts measures of social abilities in autism spectrum disorder. bioRxiv.:290320. doi:10.1101/290320.

Liaw A, Wiener M. 2002. Classification and regression by randomForest.

Marek S, Tervo-Clemmens B, Calabro FJ, Montez DF, Kay BP, Hatoum AS, Donohue MR, Foran W, Miller RL, Feczko E. 2020. Towards Reproducible Brain-Wide Association Studies. bioRxiv.

Matthews PM, Honey GD, Bullmore ET. 2006. Neuroimaging: Applications of fMRI in translational medicine and clinical practice. Nat Rev Neurosci. 7(9):732.

Noble S, Scheinost D, Constable RT. 2019. A decade of test-retest reliability of functional connectivity: A systematic review and meta-analysis. NeuroImage.:116157. doi:10.1016/j.neuroimage.2019.116157.

Noble S, Spann MN, Tokoglu F, Shen X, Constable RT, Scheinost D. 2017. Influences on the Test– Retest Reliability of Functional Connectivity MRI and its Relationship with Behavioral Utility. Cereb Cortex. 27(11):5415–5429. doi:10.1093/cercor/bhx230.

Nunnally JC. 1970. Introduction to psychological measurement.

Orru G, Pettersson-Yeo W, Marquand AF, Sartori G, Mechelli A. 2012. Using support vector machine to identify imaging biomarkers of neurological and psychiatric disease: a critical review. Neurosci Biobehav Rev. 36(4):1140–1152.

Park SH. 1981. Collinearity and optimal restrictions on regression parameters for estimating responses. Technometrics. 23(3):289–295.

Power JD, Barnes KA, Snyder AZ, Schlaggar BL, Petersen SE. 2012. Spurious but systematic correlations in functional connectivity MRI networks arise from subject motion. NeuroImage. 59(3):2142–2154. doi:10.1016/j.neuroimage.2011.10.018.

Power JD, Cohen AL, Nelson SM, Wig GS, Barnes KA, Church JA, Vogel AC, Laumann TO, Miezin FM, Schlaggar BL, et al. 2011. Functional Network Organization of the Human Brain. Neuron. 72(4):665–678. doi:10.1016/j.neuron.2011.09.006.

Reddan MC, Lindquist MA, Wager TD. 2017. Effect size estimation in neuroimaging. JAMA Psychiatry. 74(3):207–208.

Rosenberg MD, Finn ES, Scheinost D, Papademetris X, Shen X, Constable RT, Chun MM. 2016. A neuromarker of sustained attention from whole-brain functional connectivity. Nat Neurosci. 19(1):165–171. doi:10.1038/nn.4179.

Salimi-Khorshidi G, Douaud G, Beckmann CF, Glasser MF, Griffanti L, Smith SM. 2014. Automatic denoising of functional MRI data: Combining independent component analysis and hierarchical fusion of classifiers. NeuroImage. 90:449–468. doi:10.1016/j.neuroimage.2013.11.046.

Satterthwaite TD, Wolf DH, Loughead J, Ruparel K, Elliott MA, Hakonarson H, Gur RC, Gur RE. 2012. Impact of in-scanner head motion on multiple measures of functional connectivity: relevance for studies of neurodevelopment in youth. NeuroImage. 60(1):623–632. doi:10.1016/j.neuroimage.2011.12.063.

Scheinost D, Noble S, Horien C, Greene AS, Lake EM, Salehi M, Gao S, Shen X, O’Connor D, Barron DS. 2019. Ten simple rules for predictive modeling of individual differences in neuroimaging. NeuroImage.

Shehzad Z, Kelly AM, Reiss PT, Gee DG, Gotimer K, Uddin LQ, Lee SH, Margulies DS, Roy AK, Biswal BB, et al. 2009. The Resting Brain: Unconstrained yet Reliable. Cereb Cortex. doi:bhn256 [pii] 10.1093/cercor/bhn256. http://www.ncbi.nlm.nih.gov/entrez/query.fcgi?cmd=Retrieve&db=PubMed&dopt=Citation&list_uids=19221144.

Shen X, Finn ES, Scheinost D, Rosenberg MD, Chun MM, Papademetris X, Constable RT. 2017. Using connectome-based predictive modeling to predict individual behavior from brain connectivity. Nat Protoc. 12(3):506–518. doi:10.1038/nprot.2016.178.

Shou H, Eloyan A, Lee S, Zipunnikov V, Crainiceanu AN, Nebel MB, Caffo B, Lindquist MA, Crainiceanu CM. 2013. Quantifying the reliability of image replication studies: the image intraclass correlation coefficient (I2C2). Cogn Affect Behav Neurosci. 13(4):714–724.

Shrout PE, Fleiss JL. 1979. Intraclass correlations: uses in assessing rater reliability. Psychol Bull. 86(2):420–428.

Siegel JS, Mitra A, Laumann TO, Seitzman BA, Raichle M, Corbetta M, Snyder AZ. 2017. Data Quality Influences Observed Links Between Functional Connectivity and Behavior. Cereb Cortex N Y N 1991. 27(9):4492–4502. doi:10.1093/cercor/bhw253.

Smola AJ, Schölkopf B. 2004. A tutorial on support vector regression. Stat Comput. 14(3):199– 222.

Sripada C, Angstad M, Rutherford S, Taxali A, Clark DA, Greathouse T, Weigard A, Hyde L, Heitzeg M. 2020. Brain Connectivity Patterns in Children Linked to Neurocognitive Abilities. bioRxiv.

Sripada C, Angstadt M, Rutherford S. 2018 Jan 1. Towards a “Treadmill Test” for Cognition: Reliable Prediction of Intelligence From Whole-Brain Task Activation Patterns. bioRxiv. doi:10.1101/412056. http://biorxiv.org/content/early/2018/09/09/412056.abstract.

Sripada C, Angstadt M, Rutherford S, Kessler D, Kim Y, Yee M, Levina E. 2019. Basic Units of Inter-Individual Variation in Resting State Connectomes. Sci Rep. 9(1):1900. doi:10.1038/s41598-018-38406-5.

Sripada C, Angstadt M, Rutherford S, Taxali A, Shedden K. 2020. Toward a “treadmill test” for cognition: Improved prediction of general cognitive ability from the task activated brain. Hum Brain Mapp.

Sripada C, Rutherford S, Angstadt M, Thompson WK, Luciana M, Weigard A, Hyde L, Heitzeg M. 2019. Prediction of Neurocognition in Youth From Resting State fMRI. Mol Psychiatry.:1–19. doi:10.1101/495267.

Sui J, Jiang R, Bustillo J, Calhoun V. 2020. Neuroimaging-based Individualized Prediction of Cognition and Behavior for Mental Disorders and Health: Methods and Promises. Biol Psychiatry.

Termenon M, Jaillard A, Delon-Martin C, Achard S. 2016. Reliability of graph analysis of resting state fMRI using test-retest dataset from the Human Connectome Project. Neuroimage. 142:172–187.

Tian Y, Margulies DS, Breakspear M, Zalesky A. 2020. Hierarchical organization of the human subcortex unveiled with functional connectivity gradients. bioRxiv.

Van Dijk KRA, Sabuncu MR, Buckner RL. 2012. The influence of head motion on intrinsic functional connectivity MRI. NeuroImage. 59(1):431–438. doi:10.1016/j.neuroimage.2011.07.044.

Van Essen DC, Smith SM, Barch DM, Behrens TEJ, Yacoub E, Ugurbil K. 2013. The WU-Minn Human Connectome Project: An overview. NeuroImage. 80:62–79. doi:10.1016/j.neuroimage.2013.05.041.

Wager TD, Atlas LY, Lindquist MA, Roy M, Woo C-W, Kross E. 2013. An fMRI-Based Neurologic Signature of Physical Pain. N Engl J Med. 368(15):1388–1397. doi:10.1056/NEJMoa1204471.

Watanabe T, Kessler D, Scott C, Angstadt M, Sripada C. 2014. Disease Prediction based on Functional Connectomes using a Scalable and Spatially--Informed Support Vector Machine. NeuroImage. 96:183–202.

Woo C-W, Chang LJ, Lindquist MA, Wager TD. 2017. Building better biomarkers: brain models in translational neuroimaging. Nat Neurosci. 20(3):365–377. doi:10.1038/nn.4478.

Woo C-W, Schmidt L, Krishnan A, Jepma M, Roy M, Lindquist MA, Atlas LY, Wager TD. 2017. Quantifying cerebral contributions to pain beyond nociception. Nat Commun. 8:14211.

Woo C-W, Wager TD. 2016. What reliability can and cannot tell us about pain report and pain neuroimaging. Pain. 157(3):511–513.

WU-Minn HCP. 2017. 1200 Subjects Data Release Reference Manual.

Yoo K, Rosenberg MD, Hsu W-T, Zhang S, Li C-SR, Scheinost D, Constable RT, Chun MM. 2018. Connectome-based predictive modeling of attention: Comparing different functional connectivity features and prediction methods across datasets. NeuroImage. 167:11–22. doi:10.1016/j.neuroimage.2017.11.010.

Zou H, Hastie T. 2005. Regularization and variable selection via the elastic net. J R Stat Soc Ser B Stat Methodol. 67(2):301–320.

